# Rapid adaptive evolution in restored populations of a perennial plant

**DOI:** 10.64898/2026.10.27.762171

**Authors:** Lea Klepka, Hannes Wallstein, Sascha Liepelt, Norbert Hölzel, Anna Bucharova

**Affiliations:** Conservation Biology, Department of Biology, Karl-von-Frisch-Straße 8, 35043 Marburg, University Marburg; Institute of Landscape Ecology, University of Münster, Münster, Germany

**Keywords:** rapid adaptation, ecosystem restoration, phenology, hay transfer, restored population

## Abstract

Restored plant populations often face novel environmental challenges because degraded restoration sites typically differ from natural habitats. The success of ecosystem restoration may therefore depend on whether restored plant populations can adapt rapidly to these novel conditions. Yet, rapid adaptive evolution during restoration remains rarely studied.

Here, we tested for rapid adaptation in *Galium wirtgenii*, a perennial herb, within restored alluvial grasslands. Sites were restored 20 years ago by transferring hay from nearby ancient semi-natural meadows. Compared to the semi-natural donor meadows, restored sites are more productive and are mown two to three months earlier. We collected seeds from 571 mother plants from 48 restored and 18 semi-natural sites and grew their offspring in a common garden over three years. To test for differences in regeneration after mowing, we included a clipping treatment when plants were in full blossom.

Plants from restored sites flowered earlier than plants from semi-natural sites, and this difference persisted across all three years, consistent with a heritable evolutionary response to earlier mowing allowing plants at restored sites to complete reproduction before mowing. Plants from restored sites also grew more slowly in early life stages, but this effect disappeared later, suggesting maternal effects rather than genetic differentiation. We did not detect differences in regeneration after clipping, likely because regrowth rarely results in viable seeds, providing little fitness advantage.

In summary, we found evidence for rapid adaptation to novel environmental conditions in restored sites, likely contributing to successful establishment of *Galium wirtgenii* populations in the restored grasslands.

## INTRODUCTION

Environmental change, driven by shifting climatic conditions or land use change, is one of the most severe threats to biodiversity (Pörtner et al., 2021; Shaw et al., 2025; Titeux et al., 2016). Plant populations are particularly vulnerable because, as sessile organisms, they have a limited escape potential (Seth & Sebastian, 2024). However, rapid adaptive evolution may provide a mechanism by which populations can adapt to novel conditions, avoid local extinction, and persist in the long term. A key question is therefore whether plant populations can adapt rapidly enough to keep pace with ongoing environmental change (Willis et al., 2008).

Evidence for rapid evolutionary responses in plants is growing. For example, invasive plant populations often adapt rapidly to novel environmental conditions in newly colonized habitats (Akbar et al., 2025; Colautti & Barrett, 2013; Novy et al., 2013; Ziska et al., 2015). Rapid evolution has also been reported in response to climate. Experimental drought, for example, selected for earlier flowering as a drought-escape strategy (Johnson et al., 2022b), while resurrection studies showed evolutionary shifts towards earlier flowering and faster growth that are consistent with adaptation to increasing drought frequency due to climate change (Rauschkolb et al., 2023). Similarly, evolutionary change within only a few generations was demonstrated in response to changes in the biotic environment, including plant community diversity (van Moorsel, Hahl, et al., 2018; van Moorsel, Schmid, et al., 2018), pollinator communities (Ramos & Schiestl, 2019), and interactions with invasive plant species (Leger & Espeland, 2010).

One of the strong factors causing adaptive differentiation in plant populations is land use, particularly mowing and grazing (Völler et al., 2017). Frequent biomass removal can favor smaller plants with reduced above-ground investment, because this limits the fitness costs of losing biomass (Völler et al., 2013, 2017; Warwick & Briggs, 1979). Moreover, because mowing can remove reproductive structures before seed set, earlier mowing may also select for earlier flowering onset, allowing plants to complete reproduction before they are mown. In addition to escaping mowing through earlier flowering, enhanced regeneration after biomass loss might be an adaptation to frequent biomass removal, as plants can compensate for biomass loss through re-growth or reproduction (Bucharova et al., 2024; Lennartsson et al., 1998; Lin et al., 2018).

Abrupt changes in environmental conditions commonly occur when plants are introduced into degraded habitats during ecosystem restoration. For example, when restoring ecosystems on post-arable land, past land management practices such as tillage and fertilization leave degraded soils with altered water-holding capacity and nutrient availability (Brudvig et al., 2013, 2021; Klein-Raufhake et al., 2022). Restored sites may also be managed differently than natural sites, which often have a protection status (Willems et al., 2026). Consequently, restored sites often differ in vegetation composition and ecological functioning from their near-natural counterparts (Merz et al., 2026; Seabloom & van der Valk, 2003), and individual plants in restored populations commonly differ from their conspecifics in natural sites (Klein-Raufhake et al., 2022; Willems et al., 2026). Several common garden studies also demonstrated rapid evolutionary differentiation between plants in restored populations and plants from the original seed source used for restoration, although it is not fully clear if these changes are adaptive because of a lack of proper replicates (Bucharova et al., 2024; Magnoli, 2020; Magnoli & Lau, 2020).

Phenotypic differentiation among populations can arise through multiple evolutionary processes, including adaptive evolution through selection and neutral evolution through genetic drift. Distinguishing between these processes requires replication at the population level, as repeated, parallel evolutionary change across independent populations is unlikely to result from stochastic processes such as genetic drift (Kawecki & Ebert, 2004; Martínez & Lang, 2023). Past studies of evolution in restoration focused on only one or two restored populations (Bucharova et al., 2024; Magnoli, 2020; Magnoli & Lau, 2020) and thus cannot reliably identify adaptive evolution (Kawecki & Ebert, 2004). Larger studies across many restored populations that would provide more reliable evidence for rapid adaptation to novel conditions in restoration are missing so far.

Here, we used a large-scale restoration project involving populations of *Galium wirtgenii,* a perennial plant, that provides an opportunity to study rapid evolution following restoration by including multiple restored and near-natural populations as replicates. In this system, former arable land was converted into species-rich alluvial grasslands by transferring green hay, seed-containing material from remnant floodplain meadows (referred to as seminatural meadows) to restored sites (Figure 1). This restoration method has been applied to many sites within the area, creating replicated restored populations that experience similar environmental conditions.

**Figure 1:**
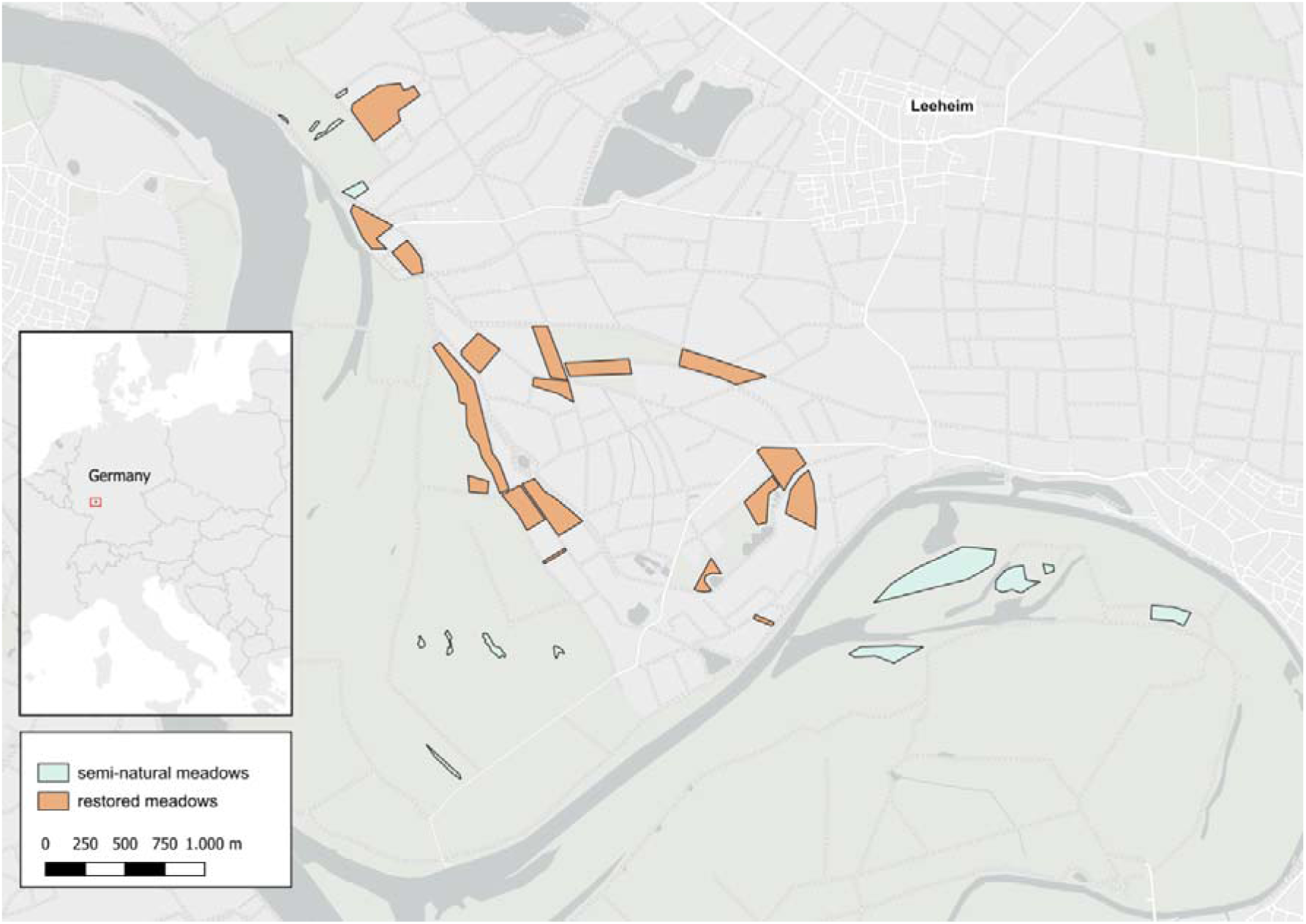
Map of the study area (Floodplain meadows in the Northern Upper Rhine Valley). Semi-natural meadows are green, restored meadows are orange.

Specifically, restored meadows are mown in mid-June, which is substantially earlier than the semi-natural meadows that are traditionally mown in August or September. Additionally, soil nutrient levels in restored meadows are higher due to previous agricultural use and fertilization. We collected seeds of *G. wirtgenii* plants from both site types and grew them in a common garden to compare flowering phenology, vegetative growth, and regeneration after mowing. To assess the potential contribution of maternal effects, we measured these traits during the first and third growing seasons. Because maternal effects often weaken over time (Weiner et al., 1997), differences that persist into later plant life are less likely the result of maternal effects. We hypothesized that earlier mowing in restored meadows has driven rapid adaptation of flowering phenology and vegetative growth. Specifically, we predicted that offspring of plants from restored meadows, compared to those from semi-natural meadows, would (1) flower earlier to complete reproduction before mowing, with this difference persisting into the third growing season; (2) invest less in vegetative growth as an adaptation to early biomass loss, with this difference also being persistent, and (3) show enhanced post-mowing regeneration.

## MATERIAL AND METHODS

### Study area

We collected seeds of our focal species, *G. wirtgenii,* from floodplain meadows in the northern Upper Rhine Valley, southwest of Frankfurt, Germany (49° 51’N, 8° 24’E). Irregular flooding by the River Rhine, followed by dry summer periods, creates variable soil moisture conditions typical of semi-natural alluvial grasslands (Šefferová Stanová et al., 2008).

The study area has undergone extensive and well-documented land-use change (Hölzel et al., 2006). Following drainage, dyking, and river regulation in the 1950s, species-rich alluvial grassland meadows were converted into arable land. From the 1980s to early 1990s, parts of the area were passively restored and managed for conservation, with mowing and grazing in autumn. However, many target species failed to re-establish, likely due to dispersal limitations (Bissels et al., 2004). Since 1997, species have been actively reintroduced using green hay transfer. Species-rich donor meadows were mown, and the hay containing diaspores of rare and endangered species was transferred to the prepared sites. Restoration outcomes have been monitored long-term (see, for example, Bissels et al., 2004; Donath et al., 2007, 2015a).

Today, semi-natural and restored meadows differ in soil conditions and management. Semi-natural sites are nutrient-poor and typically mown later in the season, whereas restored sites are still comparatively nutrient-rich and mown earlier to harvest fodder (Donath et al., 2007, 2015). The plant populations in situ also differ in traits, and across species, restored populations flower on average two days earlier than conspecifics at natural sites (Bucharova et al., 2024; Willems et al., 2026).

### Study species

We focused on *Galium wirtgenii* F.W.Schultz (Rubiaceae), a perennial characteristic of species-rich floodplain meadows in central Europe. It typically occurs in *Molinion* and *Cnidion* alliance types and flowers from May until June. *G. wirtgenii* shows a limited potential for clonal spread and relies on sexual reproduction by seeds to colonize new sites. The species is very common in most seminatural and restored meadows in the study area.

### Common garden experiment

To test for heritable adaptive change in *G. wirtgenii* populations, we conducted a common garden experiment using offspring from both sites. In June and July 2021, we collected seeds from 571 maternal individuals across 18 semi-natural and 48 restored meadows (8-10 individuals per site) and stored them at 4°C in paper bags placed in airtight containers with silica gel. Most of the sites were located within an area of 5 × 4 km, and some of the restored sites were only a few dozen meters apart. It is thus likely that there is ongoing gene flow between the sites. Before the experiment, we determined seed weight by weighing 20 seeds from each maternal plant and scaling it up to thousand-seed weight.

In February 2022, we placed 20 seeds of each maternal plant on moist filter paper (Macherey-Nagel®, MN 616 G 55 mm) into Petri dishes (Sarstedt®, 60 x 15 mm). In total, we used 20 x 571 = 11,420 seeds. We let them germinate in a climate cabinet (PHCbi® MLR-352H-PE Climate Chamber) with a 14/10 h day/night cycle at 25°C/10°C and a humidity of 65%. We germinated the seeds in Petri dishes in a climate cabinet rather than in soil in the greenhouse because the species requires warm temperatures for germination, and the available space in the climate cabinet was insufficient to germinate the seeds directly in soil. We transplanted seedlings with fully developed cotyledons into trays (QuickPot®) filled with standard potting soil (Hawita® Fruhstorfer Erde, Typ LD 80: two seedlings for a QuickPot cell, two QuickPot cells per maternal plant. After four weeks in a greenhouse (15°C), we transferred the plants in QuickPots to an outdoor common garden for acclimation and reduced the number of seedlings to one per QuickPot cell. Approximately seven weeks after germination, we transplanted the individuals into 3 L pots containing standard potting soil and arranged them in a fully randomized design in a common garden located in the botanical garden of Marburg, Germany. Plants were watered regularly throughout the experiment. In June 2022, we applied a clipping treatment to half of the plants to simulate early mowing. For this, we removed all aboveground biomass 2 cm above the soil surface in June 2022, dried it at 70°C, and weighed it.

Because of limited germination and early mortality, the final experiment comprised 99 and 208 maternal families from 11 semi-natural and 36 restored meadows, respectively. Most maternal families were represented by two plants, with one assigned to the control and one to the clipping treatment, but for some families, only one seedling was available, which we randomly assigned to one of the two treatments. In total, the final experiments contained 522 plants.

During the first season, we recorded flowering onset and biomass production. Flowering onset was defined as the date of the first open flower of an individual, calculated as the number of days from transplantation into QuickPots to first flowering. For the plants in the clipping treatment, we additionally recorded time to regeneration flowering as the number of days from biomass removal in June to the first open flower after biomass removal. We recorded early biomass production as the weight of the dry biomass from the clipping treatment. In September 2022, we cut aboveground biomass 2 cm above the soil in all plants, dried and weighed it to obtain final biomass.

In 2023 and 2024, we kept plants in the common garden without applying any treatments. Additional watering beyond natural precipitation was not necessary because the plant roots grew through the pots into the natural soil. In the third growing season (2024), we recorded flowering onset as the day of the year (DOY) for all individuals. We treated all plants identically, and when most plants started to produce seeds, we measured plant height and harvested aboveground biomass. We did not repeat the clipping treatment in 2024.

### Statistical analyses

We performed all analyses in R (R Core Team, 2026) using individual linear mixed-effects models (LMMs) for each research question and assessed the significance of fixed effects using Type III ANOVAs.

First, we tested whether there is any systematic difference in seed quality between seeds from seminatural and restored sites. To do this, we fitted an LMM with thousand-seed weight per mother plant as response variable, origin (restored vs. semi-natural meadow) as fixed factor, and seed sampling site as random factor.

Seeds from restored meadows had a significantly lower seed mass than seeds from semi-natural meadows (*P* = 0.006, R^2^ = 0.05, Table 1). To account for potential maternal provisioning effects on traits, we included seed weight as a covariate in all subsequent models.

**Table 1:**
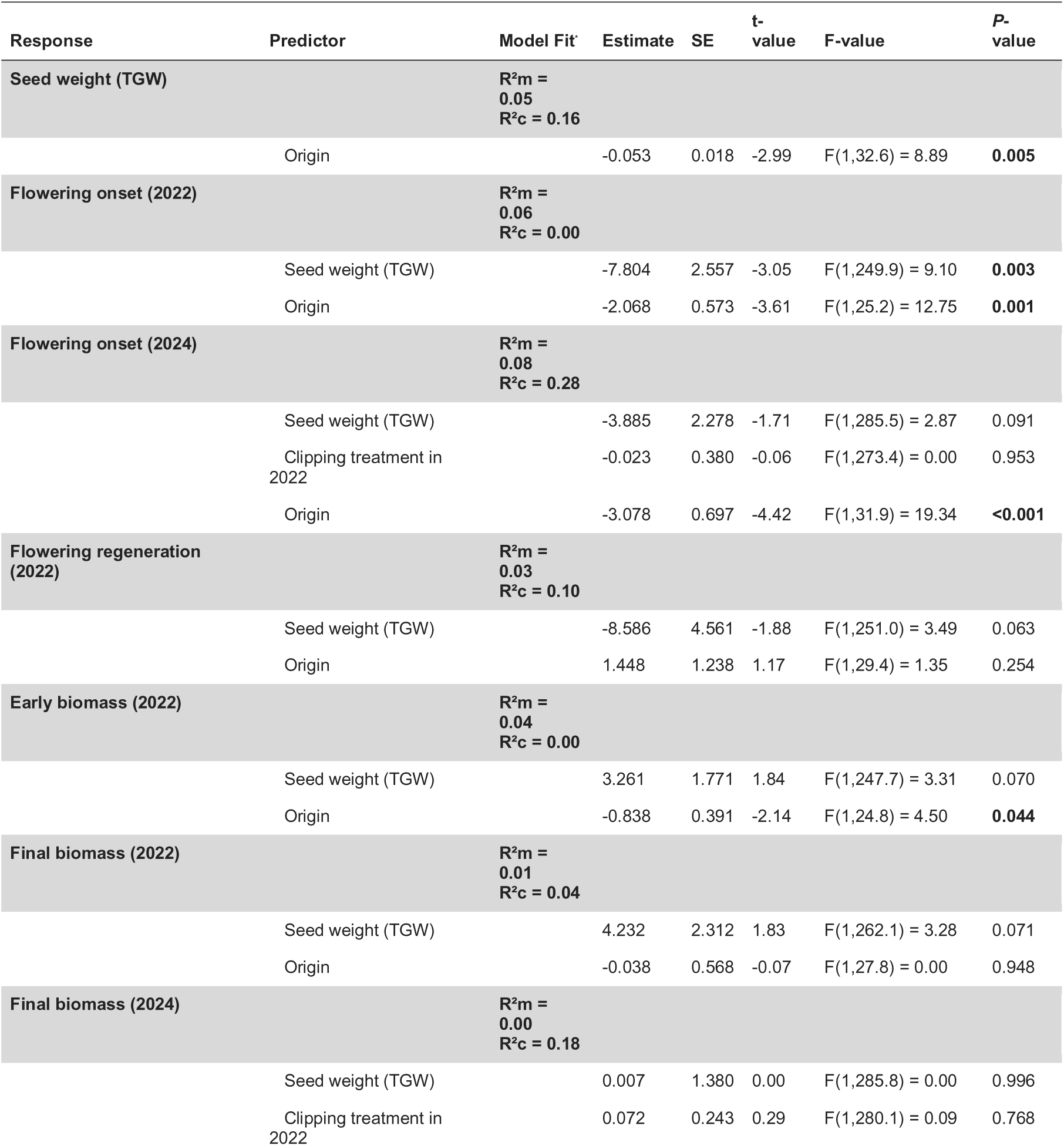
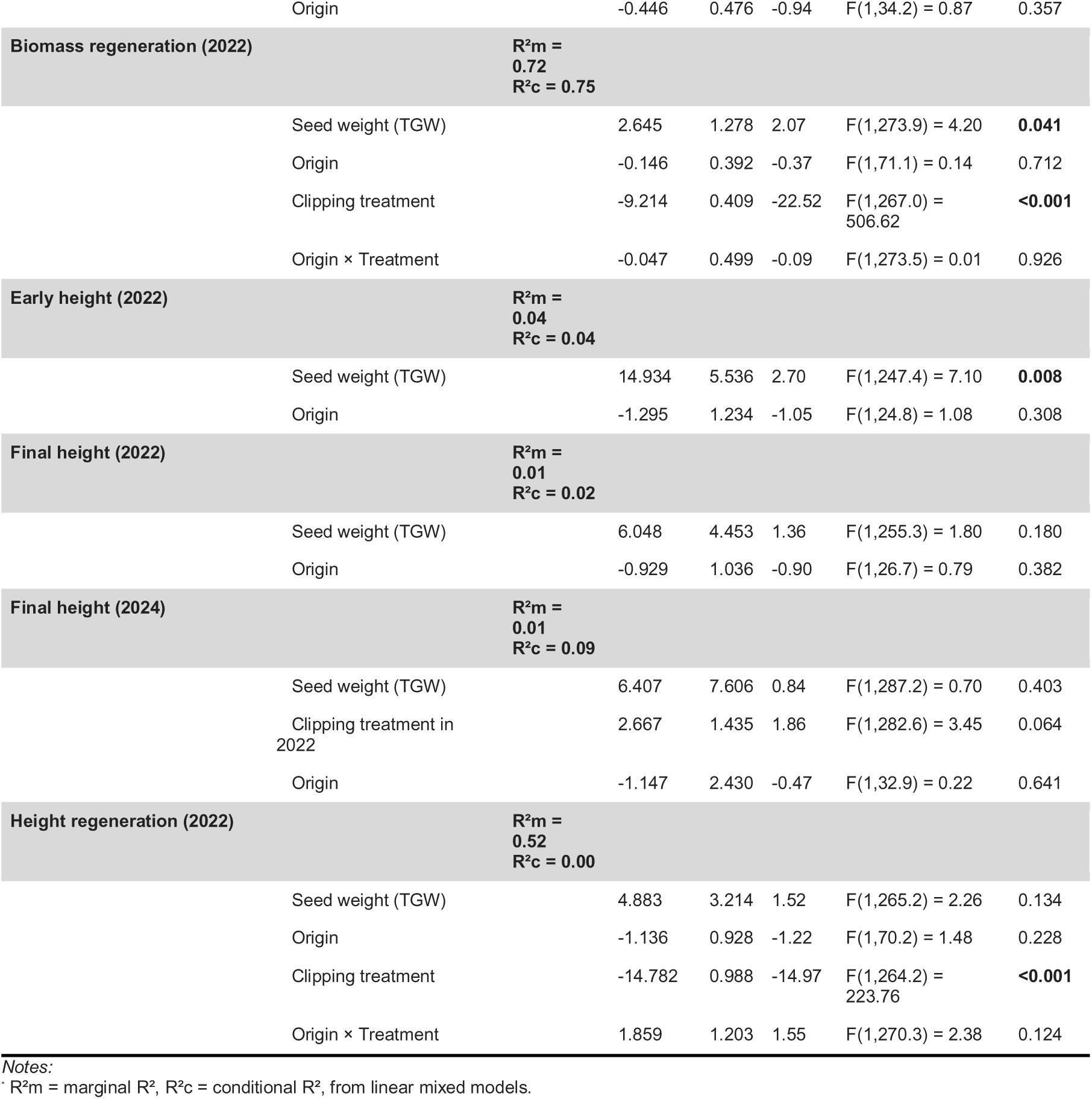
Results of linear mixed models testing the effects of seed origin (restored vs. semi-natural meadow) on flowering phenology and vegetative traits (biomass and height) in the first (2022) and third growing seasons (2024). We additionally tested differences in the thousand-grain weight between seeds originating from restored and semi-natural meadows as well as the effect of seed origin on regeneration potential following a clipping treatment in the first growing season. The estimate for Origin represents the difference from the semi-natural meadow baseline (negative values indicate earlier flowering). *P*-values are based on Type III ANOVA F-tests. Significant effects (*P* < 0.05) are highlighted in bold.

To test whether flowering phenology differed between offspring of plants from restored meadows and those from semi-natural meadows, we fitted separate models for the first and third growing seasons, because of different data structure. For the first year, we included only plants from the control treatment, as 19 individuals from the clipping treatment were harvested before flowering. The model included origin as fixed effect, seed weight as covariate, and seed sampling site as random effect. For data from the third year, we used data from all individuals because we did not apply clipping in that year. In addition to origin and seed weight, we included the first-year treatment (clipping vs. control) as covariate to account for potential long-term consequences of the clipping treatment. We also included the maternal plant identity as an additional random factor to account for relatedness among siblings.

We tested for differences in vegetative growth using above-ground biomass and plant height as response variables. In the first growing season, we analyzed two time points separately: first, to assess early growth, we used biomass from individuals harvested during the clipping treatment; second, to assess final growth, we used data from control individuals harvested at the end of the first growing season. Both models included origin as fixed factor, seed weight as covariate, and seed sampling site as random effect. For data from the third growing season, when we did not apply clipping, we fitted LMMs with third-year biomass and height as response variables for all plants. We included origin, seed weight, and first-year treatment as fixed factors, and both seed sampling site and the maternal plant as random factors.

Lastly, we assessed post-mowing regeneration. For flowering time after clipping, we used only data from individuals in the clipping treatment and fitted an LMM with origin as fixed factor, seed weight as covariate, and seed sampling site as random factor.

For final biomass and plant height following biomass removal, we used data from all individuals from both treatments and fitted LMMs with origin, treatment, and their interaction as fixed factors, seed weight as a covariate, and seed sampling site as well as maternal plant as random factors. In all models, we verified model assumptions by visual inspection of residual plots and QQ plots following Zuur et al. (2010) and used LME4, LMERTEST, and CAR-packages for LMMs (Bates et al., 2015; Fox et al., 2024; Kuznetsova et al., 2017), the MUMIN package for calculating approximate conditional R^2^ (Bartoń, 2022), and GGPLOT2 for visualization (Wickham, 2016).

## RESULTS

We found evidence for differentiation in flowering phenology and early vegetative growth between restored and semi-natural populations of *G. wirtgenii* (Table1, Figure 2). Offspring from restored meadows flowered 2 days earlier than those from semi-natural meadows in the first growing season. This differentiation persisted into the third growing season and was even stronger (flowering onset 3 days apart, Table 1, Figure 2A). The effect of origin on vegetative growth was only weak: Offspring from restored meadows had slightly lower early biomass, but biomass at the end of both the first and the third growing seasons did not differ between origins (Table 1, Figure 2B). Plant height did not differ at any assessed time point (Table 1, Figure 2C).

**Figure 2:**
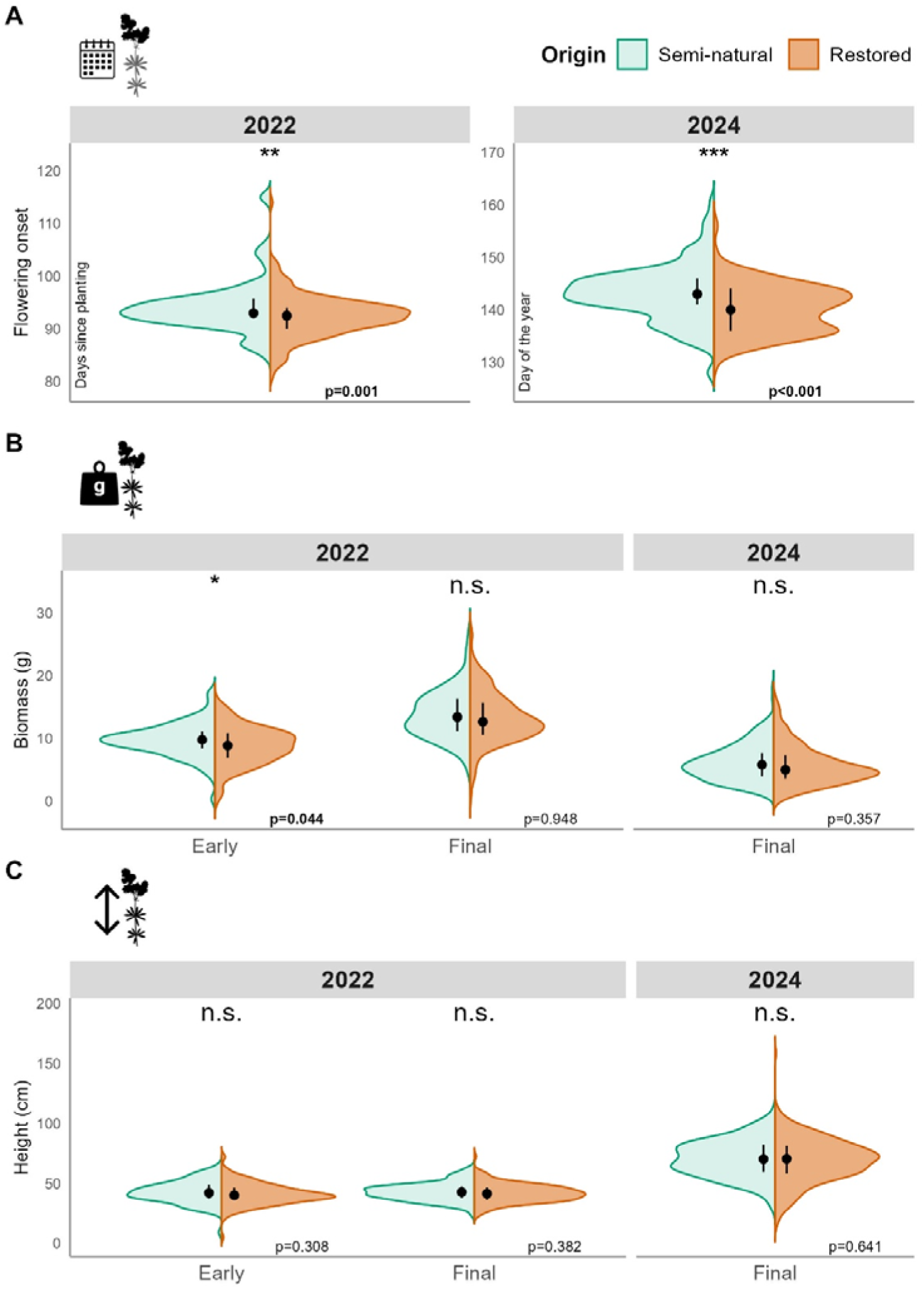
Effects of seed origin (semi-natural vs. restored meadows) on flowering phenology and vegetative traits measured in 2022 and 2024. (A) Flowering onset, shown as days since planting (2022) and day of the year (2024). (B) Aboveground biomass and (C) plant height, each shown separately for early and late harvest in 2022 and late harvest only in 2024. The black points represent the median with the interquartile range (25^th^ – 75^th^ percentile). Asterisks indicate the significance of the origin effect. See Table 1 for individual model results.

Offspring from restored meadows showed no enhanced post-mowing regeneration (Table 1, Figure S1). They did not re-flower earlier, nor did they grow taller or produce more biomass after mowing compared to plants from semi-natural meadows.

## DISCUSSION

Plant populations that establish in degraded habitats within restoration projects face novel and often challenging environmental conditions. To persist, plants may cope with these novel conditions through phenotypic plasticity. Alternatively, such conditions can impose selective pressures that shift heritable trait variation toward phenotypes that are better suited for these conditions. Such a process can result in rapid adaptation and can contribute to restoration success. Here we show, using a common garden experiment with a perennial *Galium wirtgenii*, that flowering phenology at restored sites shifted by more than two days in comparison to the near-natural sites that served as seed source for the restoration. Given the large number of replicate sites in both restored and semi-natural sites, it is unlikely that this pattern results from neutral evolution (Kawecki & Ebert, 2004; Martínez & Lang, 2023). Instead, the consistent difference in flowering phenology strongly suggests adaptive evolution, likely driven by the altered mowing regime. In line with recent findings (Bucharova et al., 2024; Magnoli, 2020; Magnoli & Lau, 2020), our results indicate that rapid adaptation to novel conditions in restored ecosystems may be common.

The earlier flowering of plants from restored meadows is likely advantageous under the early mowing regime because it increases the probability of successful reproduction before biomass removal. Since mowing removes reproductive structures during the flowering and seed ripening period of *G. wirtgenii*, this regime is likely to favour individuals that flower earlier, increasing the probability of completing reproduction and producing mature seeds before biomass removal (Willems et al., 2026). In contrast, mowing on semi-natural meadows occurs later in the year, after seed set in this species, and therefore does not represent a selective pressure on flowering time. Thus, early mowing shortens the reproduction period and may favour rapid completion of the reproductive cycle. This pattern is in line with studies showing that early or frequent mowing is generally selective on flowering phenology across grassland species (Völler et al., 2013, 2017). Similar shifts towards earlier flowering have been reported in response to drought (Franks et al., 2007; Johnson et al., 2022; Kulpa & Leger, 2012; Villhauer et al., 2026) and shorter growing seasons along climatic gradients (Colautti & Barrett, 2013; Novy et al., 2013). In summary, these findings support the ecological plausibility of selection for advanced flowering under shortened vegetation periods, in our case caused by earlier mowing.

The consistent and persistent difference in the flowering onset across multiple populations under common conditions provides strong evidence for adaptive rather than neutral evolution. In fact, earlier flowering in restored meadows in the same study system has been shown in situ across 16 species (Willems et al., 2026). Bucharova et al. (2024) reported such a flowering advancement in a common garden for one pair of restored and seminatural meadow, but the study design included only one replicate and, therefore, could not tease apart adaptive evolution and genetic drift. Similarly, other studies on rapid evolution in restoration focused on a maximum of one or two populations and provided ambiguous results (Magnoli, 2020; Magnoli & Lau, 2020). Because the current study included replicated restored and near-natural sites, it provides the strongest evidence so far for adaptive evolution in restored plant populations.

The observed differences between plants from restored and near-natural populations could be caused by maternal environmental effects, but this is rather unlikely. We observed the difference in phenology even in three-year-old plants, while maternal effects mainly affect early life stages and weaken as the plants grow older (Weiner et al., 1997) particularly for reproductive traits (Bischoff & Müller-Schärer, 2010).

Additionally, we observed the main differentiation in flowering time, a trait that is rather genetically determined than environmentally driven (Villhauer et al., 2026). Although a “refresher generation” under common conditions would be required to fully exclude maternal effects, it is more likely that the differentiation we observe is heritable and thus the result of selection on standing genetic variation.

Plants from restored meadows did not show enhanced regeneration after the clipping treatment. They neither started to flower earlier nor showed faster vegetative regrowth after clipping. Previous studies have shown differentiation in regrowth after biomass removal in other species, suggesting that responses to mowing and grazing can differ between populations (Carman & Briske, 1985; Kirschbaum et al., 2021).

However, these responses vary among species and management regimes, with interannual variation likely being more important than mean mowing or grazing intensity (Kirschbaum et al., 2021). In our study system, the hot and dry summers may limit quick regeneration after mowing, potentially reducing the advantage of fast regrowth. However, we applied the clipping treatment only in the first and not the third growing season, and we cannot completely exclude differences in later life stages.

We found no consistent differentiation between restored and semi-natural populations in vegetative growth, measured as aboveground biomass and plant height. Offspring plants from restored meadows had slightly lower biomass early in the first growing season, but final biomass did not differ either in the first or the third season, which is in line with previous findings of Bucharova et al. (2024). Seeds collected in semi-natural meadows were larger, which could potentially enhance germination and early growth through greater nutrient reserves (Vaughton & Ramsey, 2001). However, because we have statistically accounted for seed mass, this alone cannot explain the observed early biomass difference. Maternal effects can also influence the offspring phenotype through seed-mass-independent mechanisms (Zas et al., 2013), a mechanism that we believe is the most likely explanation. This is supported by the observation that differences were present only in early life stages and absent by the end of the first growing season, which is typical of maternal effects (Bischoff & Müller-Schärer, 2010). Thus, the lack of a persistent biomass difference suggests that the observed growth patterns do not reflect genetic differentiation between populations from semi-natural and restored meadows.

### Implications for restoration

We demonstrate that restored plant populations can show adaptive evolutionary changes in response to land-use management within only two decades. This finding, together with previous studies documenting rapid evolution in individual restored populations, indicates that rapid adaptation in restoration does happen and possibly positively influences restoration success. This shows that any possible pre-adaptation of the seed material for restoration is further modulated after the restored populations have established.

Our findings may also have consequences for selecting the optimal seed provenance for restoration. In many regions, practitioners strictly require local sourcing of the seeds because of local adaptation (Kaulfuß et al., 2022). In the case of our study system, the plant material was sourced from local near-natural sites. However, the evolved phenology difference between restored and natural populations (2 days) is in the same order of magnitude as differences observed among provenances that would be sourced from 200-300 kilometers away or from climates that differ by 1-2°C from the local climate (Bucharova et al., 2017), as well as flowering advancements associated with rising temperatures (3.6 days per 1°C warming, (Willems et al.,

2022). Given that such differences evolved over just two decades, it is possible that local adaptation might be modified anyway by rapid evolution on site and provenances from a broader region and climate similarity would perform equally well. In this context, strict adherence to local provenance might not be essential. Instead, practitioners may prefer sourcing seeds from multiple populations from the broader region, as this could provide greater genetic variability for selection to act on and thereby facilitate rapid adaptation to local conditions. This may be particularly relevant under ongoing climate change, where high genetic variation could increase the adaptive potential of restored populations to respond to changing climatic conditions.

## Author contribution

L.K.: investigation, formal analysis, data curation, writing original draft, visualization H.W.: investigation, review and editing. S.L.: supervision, review and editing. N.H.: conceptualization, review and editing. A.B.: conceptualization, methodology, resources, writing original draft, review and editing, supervision

## Acknowledgement

We thank Christina Mengel for technical support.

## Data Availability Statement

The data supporting the findings of this study are not publicly available at the time of submission but will be deposited in an appropriate public repository upon acceptance of the manuscript. Data will be made available to editors and reviewers upon reasonable request.

---

**Figure S1:**
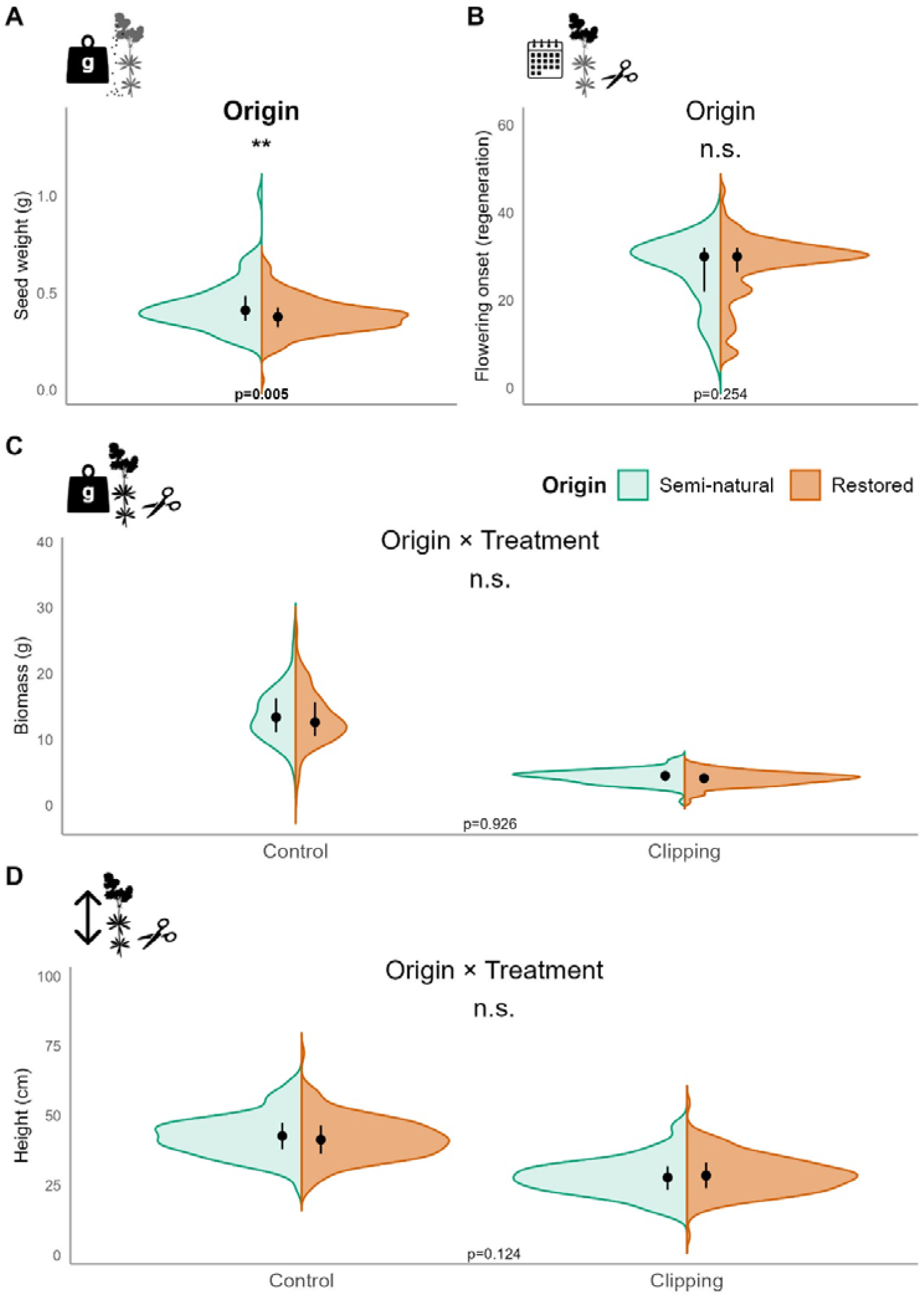
Effects of seed origin (semi-natural vs. restored meadows) on seed traits and the regeneration after mowing. (A) Seed weight of field-collected seeds. (B) Second flowering onset, shown as days since mowing. (B) Aboveground biomass and (C) plant height. The black points represent the median with the interquartile range (25^th^ – 75^th^ percentile). Asterisks in (A) and (B) indicate the significance of the origin effect; asterisks in (C) and (D) indicate the significance of the origin × treatment effect. See Table 1 for individual model results.

